# Spatiotemporal functional organization of excitatory synaptic inputs onto macaque V1 neurons

**DOI:** 10.1101/558163

**Authors:** Niansheng Ju, Yang Li, Fang Liu, Hongfei Jiang, Stephen L. Macknik, Susana Martinez-Conde, Shiming Tang

## Abstract

The integration of synaptic inputs onto dendrites provides the basis for computation within individual neurons. Whereas recent studies have begun to outline the spatial organization of synaptic inputs on individual neurons, the underlying principles related to the specific neural functions is not well known. Here we performed two-photon dendritic imaging with genetically-encoded glutamate sensor in awake monkeys, and successfully mapped the excitatory synaptic inputs on dendrites of individual V1 neurons with high spatial and temporal resolution. We found that although synaptic inputs on dendrites were functionally clustered by feature, they were highly scattered in multidimensional feature space, providing a potential substrate of local feature integration on dendritic branches. We also found that nearly all individual neurons received both abundant orientation-selective and color-selective inputs. Furthermore, we found apical dendrites received more diverse inputs than basal dendrites, with larger receptive fields, and relatively longer response latencies, suggesting a specific apical role in integrating feedback in visual information processing.

---

Sensory information processing requires that cortical neurons sample from their dendritic synaptic inputs to serve as the basic unit of computation^1^. Whereas *in vitro* patch-clamp and intracellular recordings have advanced our understanding of how dendritic synaptic integration occurs in single neurons^2,3^, and *in vivo* studies have extended those methods to highlight how dendritic activity contributes to cortical functions such as orientation and direction selectivity^4,5^, sample size and spatial resolution have limited the utility of these methods. High-resolution two-photon calcium imaging of dendrites, in contrast, can achieve long-term functional mapping of individual dendritic inputs in the intact brain ^6–13^. Additionally, determining the temporal sequencing between dendritic inputs will also render new and critical insights^14,15^. A recently developed genetically encoded glutamate-sensing fluorescent reporter, iGluSnFR^16^, which has a high signal-to-noise ratio (SNR) and fast kinetics—promises to map spatiotemporal functional organization of dendritic excitatory inputs.

In this study, we performed two-photon dendritic imaging with iGluSnFR in awake macaque monkeys, and obtained fine functional spatiotemporal maps of dendritic excitatory inputs in individual V1 neurons.

## Results

### Measuring excitatory synaptic inputs onto dendrites of macaque V1 neurons

We performed two-photon imaging on dendritic shafts of individual V1 neurons expressing SF-iGluSnFR.A184S (**Fig. 1a** and **2a**) while monkeys fixated and viewed visual stimuli. Robust and spatially localized fluorescence increases were evoked on dendrites, which we refer to as regions of interest (ROIs) (see Methods for details of analysis, **Fig. 1b**). We found that even adjacent ROIs could possess a wide variety of preferences between and within domains of orientation, spatial frequency (SF) and color (**Fig. 1c, d**, **Fig. 2b-d**, and **Supplementary Fig. 1**). Responses of individual ROIs exhibited a high-degree of stability across trials (mean responses to tuned stimuli ± STD = 0.73 ± 0.12, whereas untuned responses exhibited negligible activity = 0.03 ± 0.29; **Fig. 1e, f** and **Supplementary Fig. 2**). Across the 1813 dendritic ROIs obtained from 3 monkeys (from 23 neurons, with 79 ± 22 dendritic ROIs per neuron), the mean ROI response intensity was remarkably uniform per monkey (0.28 ± 0.03, 0.29 ± 0.03 and 0.23 ± 0.04 for each monkey, mean ± STD; **Fig. 1g**). Thus high quality dendritic imaging of glutamate responses in awake macaque monkeys enables precise determination of the functional properties of excitatory synaptic inputs.

**Fig. 1.**
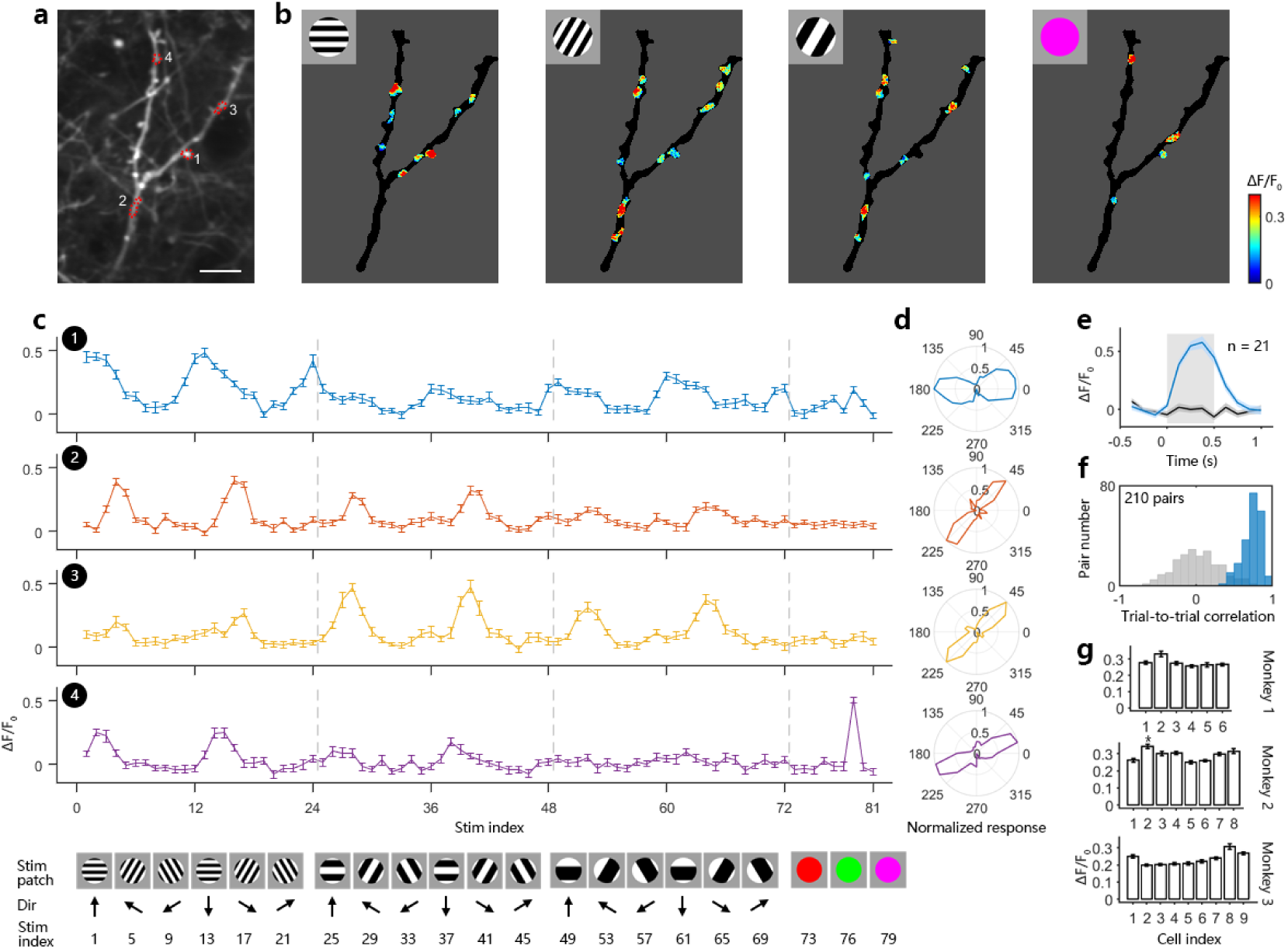
Imaging excitatory inputs on dendritic shafts in awake macaque monkey V1. **a**, Dendritic shafts of one neuron expressing SF-iGluSnFR.A184S in superficial layer of V1. Red dotted circles are representative ROIs; scale bar, 10 µm. **b**, Fluorescent responses on dendritic shafts elicited by visual stimuli (insets). Intensity of iGluSnFR fluorescence signals (of greater than 2.5 STD over baseline) are denoted by the colorbar (right) **c**, Responses from the numbered ROIs from panel **a**, plotted as a function of stimulus (below; error bar, SEM). Top, response curves; bottom, a subset from the entire set of 81 visual stimuli consisting of either 1 of 9 different color patches, or a drifting grating having 1 of 12 orientations, with 1 of 2 drift directions, and 1 of 3 spatial frequencies. **d**, Orientation polar plots from the optimal SF for each ROI. **e**, Fluorescence traces of ROI 1 corresponding to the highest (blue) and lowest (black) response intensity from all stimuli tested. Curve shadow, SEM; square shadow, stimuli onset interval. **f**, Repeatability of worst (gray) and best (blue) stimulus responses across trials as in **e** were calculated as trial-to-trial correlations. **g**, Mean value of the optimal response to each of the 1813 ROIs averaged with each the 23 source neurons (presented separately by monkey; error bar, SEM; asterisk indicates significant difference when comparing to the first column (Kruskal-Wallis test and a multiple comparisons of the group means; p < 0.05).

**Fig. 2.**
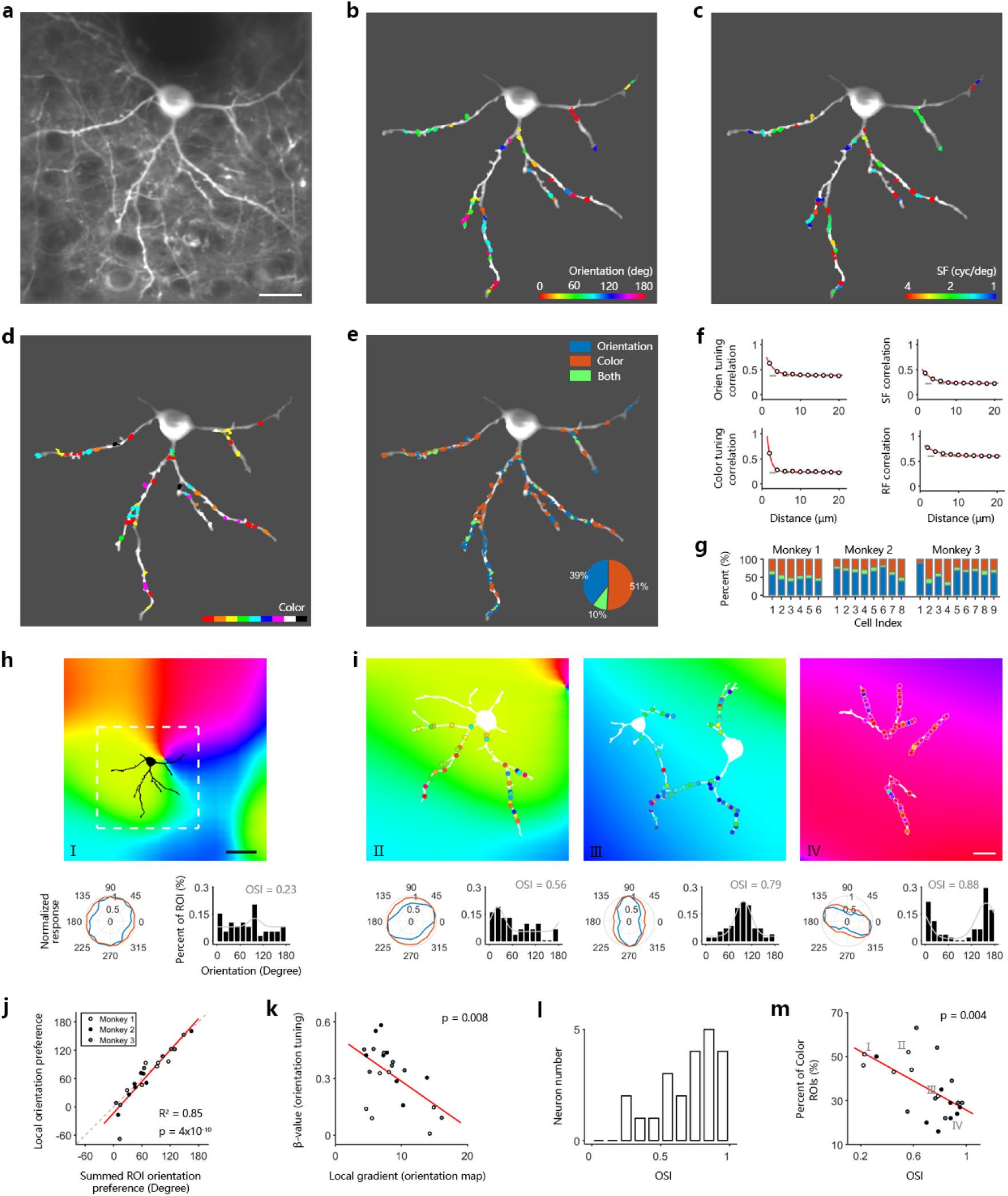
The spatial organization of dendritic excitatory inputs on individual V1 neurons. **a**, Two-photon image of an example neuron expressing SF-iGluSnFR.A184S in V1 superficial layers. Scale bar, 20 µm. **b**, Map of orientation-selective ROIs on dendrites of the example neuron. Orientation preferences are colored for vector sum polarity for each ROI (refer to colorbar at the lower right). **c**, Map of SF preferences of the orientation-selective inputs. SF preferences are colored for fitting maximum for each ROI (refer to colorbar at the lower right). **d**, Map of color-selective inputs. Color preferences are labeled with the dominant polarity for each ROI. **e**, A mixture map of oriented and colored inputs. The inserted polar plot denotes the proportion of orientation versus color inputs (ROI areas). **f**, Relationship between the dendritic distance of ROI pairs (10956 ROI pairs in total from 23 neurons) and their tuning correlation coefficients (top left, orientation tuning correlation among orientation-selective ROIs; top right, SF tuning correlation among orientation-selective ROIs; bottom left, color tuning correlation among color-selective ROIs; bottom right, RF correlation among orientation-selective ROIs). Data points are fitted with an exponential curve (red), and shuffled data are presented as a reference (gray dashed). Error bar, SEM. **g**, Proportion of orientation inputs versus color inputs (ROI area) for all 23 measured neurons. **h**, Top, neuron position on local orientation map. White dashed square, 166 µm in width and height, corresponding to 100 pixels in raw two-photon images; scale bar, 50 µm. Bottom, left, normalized polar plots of orientation tuning of summed ROI responses (blue lines) and local cortical orientation selectivity (averaged response within a local area marked by the white dashed square; red lines); right, frequency distribution of ROIs’ preferred orientation fitted with Gaussian functions (grey curves). OSI, orientation selectivity index. **i**, Dendritic orientation map of single neurons overlaid on local population orientation maps. Scale bar, 20 µm. **j**, Local cortical orientation preference versus that of summed dendritic ROI responses for each single neuron. Red line, regression line; grey dashed line, unity line. R^2^ = 0.85 when fitted to the unity line. **k**, β-value of dendritic orientation tuning versus local gradient of orientation preference. **l**, Histogram of neurons with varying OSI. **m**, Proportions of color-selective ROIs for each neuron versus its OSI.

### Synaptic inputs are moderately clustered by feature

We first examined how the excitatory synaptic inputs on dendritic shafts are clustered for each single visual property tested. Consistent with findings in anesthetized lower mammals, we found moderate clustering within local dendrites along single feature dimensions of receptive field (RF) and orientation (λ = 3.4 μm and 2.0 μm respectively, see Methods for details of analysis; **Fig. 2f**), as well as SF and color (λ = 2.5 μm and 0.9 μm). We also found that the orientation of dendritic inputs to individual neurons tended to match local orientation columnar maps (p < 10^−9^, linear regression; **Fig. 2h-j**). Input preferences varied in consistency as a function of the cortical column orientation preference map gradient (p < 0.01, linear regression; **Fig. 2k**).

### Functional trade-off between orientation/color inputs in individual V1 neurons

Interestingly, nearly all individual neurons received both orientation-selective and color-selective dendritic inputs (46% vs 45%, 63% vs 28% and 58% vs 33% for 3 Monkeys respectively; **Fig. 2e, g**). Most inputs tended to be either oriented or colored, and were interdigitated on the individual dendrites (AUC of ROC = 0.46 ± 0.08, mean ± STD across 23 neurons from 3 monkeys; **Fig. 2e, g** and **Supplementary Fig. 3**). The proportion of orientation/color mixing varied across individual neurons (±7%, ±12% and ±19% oriented inputs for each monkey, ±STD; **Fig. 2g**). Input homogeneity correlated with the proportion of each input type for each neuron. Neurons that had relatively larger amount of orientation-selective inputs tended to also receive homogenous orientation-selective synaptic inputs (p < 0.005, linear regression; **Fig. 2l, m**), and vice-versa for color-selective inputs (p < 0.01, linear regression; **Supplementary Fig. 4**). This orientation/color trade-off suggests a functional competition between oriented and colored inputs within individual V1 neurons.

**Fig. 3.**
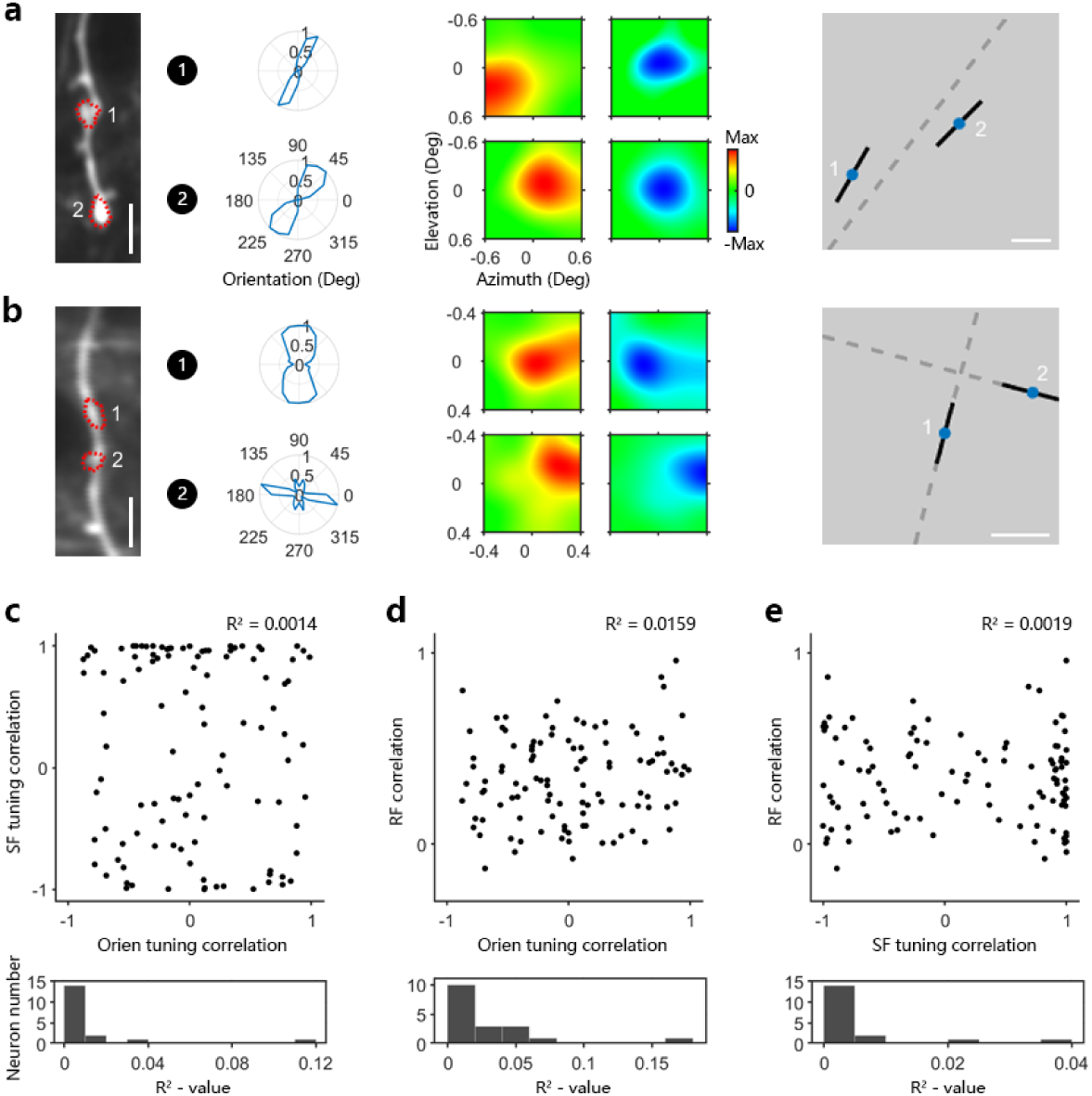
Diversity of dendritic inputs among multi-feature dimensions. **a**, Two ROIs on one dendritic draft share similar orientation preferences while having distinct RFs. Left, sample dendrite and target ROIs. Scale bar, 5 µm. Middle, the corresponding orientation preferences and RFs (red, ON; blue, OFF). Right, preferred orientation (black) overlaid at the site of each RF center (blue) for each ROI. Scale bar, 0.2 degree. **b**, Another sample dendrite with an associated ROI pair. Those ROIs have both different orientation preferences and different RFs. **c**, Inter-ROI SF tuning correlation as a function of orientation tuning correlation. Top, scatter plot from a sample neuron; R-square value from linear regression. Bottom, distribution of R-square value from all collected neurons. **d**, Inter-ROI RF correlation versus orientation tuning correlation. **e**, Inter-ROI RF correlation versus SF tuning correlation.

**Fig. 4.**
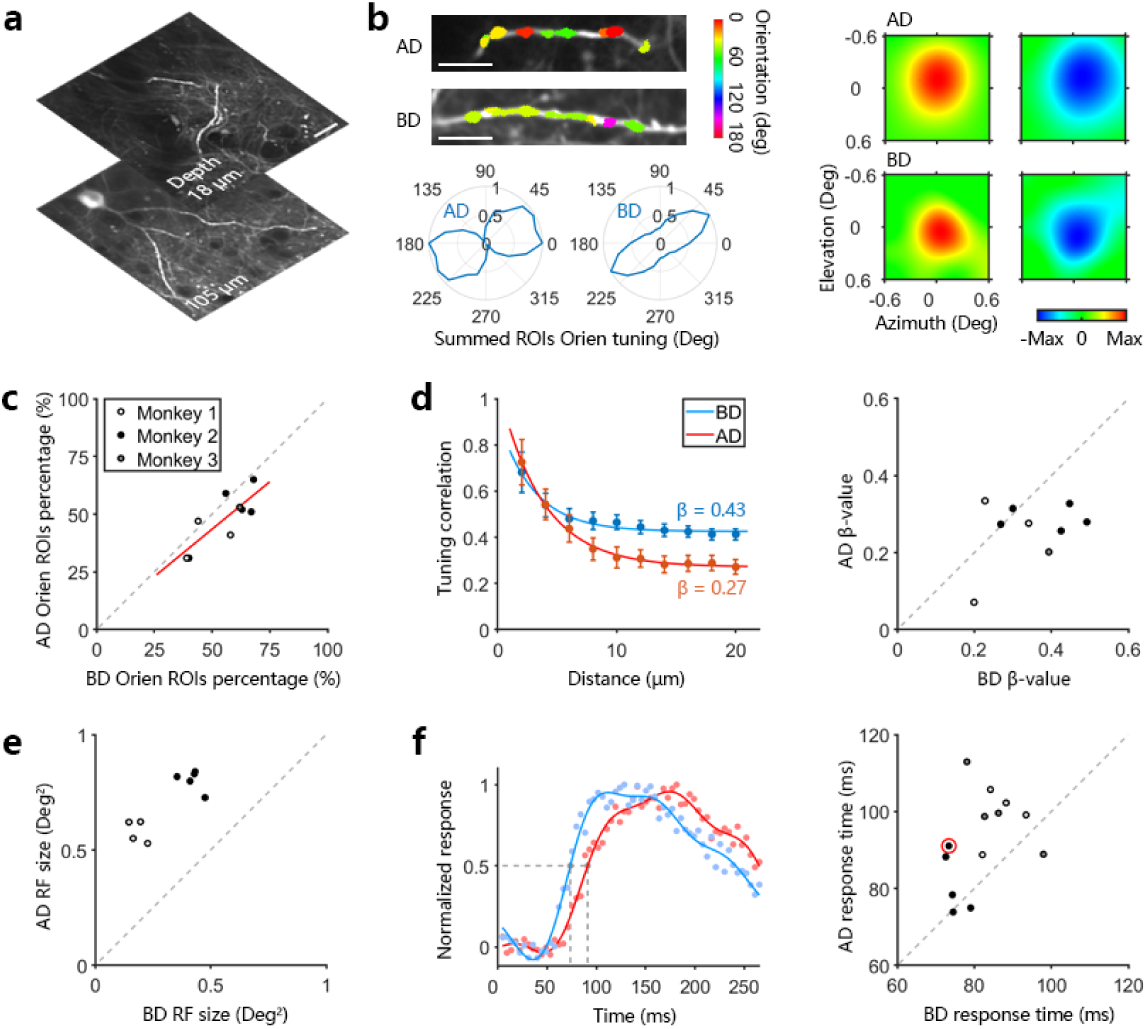
Comparison of excitatory dendritic inputs on apical dendrites versus those on basal dendrites. **a**, Apical and basal dendritic shafts of one example neuron. Scale bar, 20 µm. **b**, Orientation preference and RF of dendritic inputs on AD and BD. Top left, orientation preferences of ROIs on AD and BD respectively, colored for vector sum polarity for each ROI. Lower left, summed orientation preference of all orientation selective ROIs on AD and BD respectively (AD, 21 ROIs on 2 dendrites; BD, 43 ROIs on 3 dendrites). Right, example RFs of two ROIs on AD and BD respectively. **c**, Proportions of orientation selective ROIs versus color selective ROIs on AD and BD of each neuron. Red line, linear fit (p = 0.009); gray dashed line, unity line. **d**, Left, relationship between the dendritic distance of ROI pairs and their tuning correlation coefficients on AD (red) and BD (blue) of the example neuron. Error bar, SEM. Right, AD β-value versus BD’s of 9 neurons (neurons with less than 15 ROIs on it’s AD were discarded). **e**. AD RF size versus BD’s of each neuron. **f**. Time courses of response on AD versus BD. Left, normalized fluorescence traces on AD (red) and BD (blue) of the example neuron. Dots, raw data points; lines, Fourier fit; grey dashed lines, marking response time points according to half-height. Right, AD response time versus BD’s of each neuron.

### Synaptic inputs are highly scattered in multidimensional feature space

We found each individual synaptic input could have a unique combination of feature preferences (including orientation, SF and RF; **Fig. 1c and Fig. 3a, b**), often scattered across the multidimensional feature space. One local pair of synaptic inputs might share similar orientation preference while possessing quite dissimilar RFs (**Fig. 3a**). In contrast, another pair of synaptic inputs might have both different orientations and dissimilar RFs (**Fig. 3b**), or different orientations with similar RFs. To quantify this scatter of features, we plotted the pairwise correlation between each input pair’s preference for orientation, SF and RF. We found that the pairwise correlations across different features was highly scattered (mean *R*^2^ = 0.01, *R*^2^ = 0.03 and *R*^2^ = 0.01 for SF versus orientation, RF versus orientation and RF versus SF respectively; **Fig. 3c-e**). In some cases, synaptic inputs onto an individual neuron were quite homogenous within a feature dimension, but quite heterogeneous in other feature dimensions (**Supplementary Fig. 5**). This wide scattering served to maximize the pool of potential matches between dissimilar features within local dendritic branches (see **Fig. 3a, b**). As such, this provides a potential computational substrate for multidimensional feature integration at the dendritic level in V1 superficial layer neurons.

### Functional distinction between apical and basal dendrites

Excitatory synaptic inputs onto apical dendritic tufts, which contain a mass of convergent long-range cortical and subcortical projections^17,18^, have been assumed essential for receiving feedback from higher level brain regions ^19,20^. Recent *in-vivo* experiments combining two-photon calcium imaging and intrinsic optical imaging have provided a functional comparison between V1 neuronal population and feedback axonal projections^21,22^. Nevertheless, a direct comparison of functional inputs to apical dendrites versus basal dendrites in single units has never been reported. We compared apical inputs versus basal inputs in individual neurons, with respect to their functional preferences and response latencies (**Fig. 4a**). Apical and basal dendrites received a similar proportion of orientation-selective versus color-selective inputs (48 ± 12% oriented inputs on AD, 55 ± 11% on BD, mean ± STD; p = 0.21, Wilcoxon Rank Sum Test; **Fig. 4c**). Their respective summed orientation preferences were also similar (*R*^2^ = 0.81 fit to the unity line; **Supplementary Fig. 6**). Apical dendrites were nevertheless significantly more diverse in function (β-value pairwise preference correlations: 0.26 ± 0.08 on AD vs 0.34 ± 0.10 on BD, mean ± STD; p = 0.047, cell-by-cell comparison paired t-test; **Fig. 4b, d**). Consistent with larger RFs previously observed feedback boutons to layer 1 of V1^21^, we found that the mean RF sizes of apical inputs were significantly larger than for basal inputs (1.41 ± 0.25 square degree on AD vs 0.63 ± 0.26 on BD; p = 2 × 10^−7^, paired t-test; **Fig. 4b, e**). We acquired precise latencies for AD vs BD inputs by imaging small regions at 226 Hz (68 × 68 μm with 64 × 64 pixel resolution), and found that apical input latencies were on average 10 ms higher than in basal inputs (92 ± 12 ms for AD and 82 ± 8 ms for BD, mean ± STD; p = 0.008, paired t-test; **Fig. 4f**). The combination of larger RF sizes, increased diversity of inputs, and higher response latency indicates that feedback dominates apical inputs.

## Discussion

Our findings provide a novel approach to identify the spatiotemporal organizational principles for excitatory dendritic inputs to V1 superficial neurons of awake macaque monkeys. Whereas dendritic imaging techniques were previously applied in anesthetized lower mammals with calcium indicators ^5–13^, no prior research had ruled out the potentially significant effects of anesthesia on spontaneous neuronal firing rates, the level of inhibition and dendritic membrane properties^1^, while isolating purely excitatory inputs onto dendrites. Critically, no mammals other than old world primates have homologous visual processing capabilities to those of human beings, with respect to color, acuity, and visual cognition ^23^. Our use of the glutamate sensor iGluSnFR removed a technical hurdle that was extant in previous studies, by eliminating the need to post-process our data to control for the effects of back-propagating spikes, which are a source of noise in Ca-imaging experiments. In our hands iGluSnFR resulted in strong signals on neuronal dendrites, whereas dendritic Ca-imaging in dendrites largely failed to provide strong signals in macaque monkeys (**Supplementary Fig. 8**). Note that somal imaging of GCaMP6s Ca-indicators nevertheless succeeds in macaque V1^24^. One possible explanation for this discrepancy is that macaques have relatively reduced ion channel densities in V1 pyramidal dendrites^25^. It follows that high-quality afferent Ca-signals will be difficult to record with high a signal/noise ratio. This interspecies difference highlights the potentially different fundamental computational functions of dendrites in primates versus those in lower mammals.

Recent two-photon dendritic imaging studies have revealed that, despite the understandable excitement in the field concerning the discovery of functional input clustering^11,12,26,27^, highly scattered distributions of inputs also commonly exist^6–8,11,13^. There is no known explanation for this difference, but combined with our results in macaques, the evidence suggests that dendrites on visual neurons that reside within brains having orientation maps that receive functionally clustered synaptic inputs on their dendrites^12^, whereas in brains without an orientation structure, synaptic inputs cluster less^6,8,11^.

Given that neuronal recordings in V1 typically find that cells are either orientation- or color-selective, our finding of significantly mixed orientation and color inputs that functionally trade-off in V1 neurons is unexpected. Critically, the higher the proportion of orientation-selective inputs a given neuron received, the more uniform those inputs tended to be, which presumably contributes to the provenance of orientation selectivity in V1 neurons^9,10^. We also found that dendritic inputs having a wide scattering of functional properties within the multidimensional feature space of the visual system potentially leads to integration both within and across visual domains, which may provide a dendritic computational mechanism for multidimensional feature integration in V1. Our findings further indicate that synaptic inputs on apical dendrites reflect feedback within the visual hierarchy^21,22^, bring new insight into function of the most iconic feature of the cortical pyramidal cell. These functional observations, in combination with the first successful application of a newly developed glutamate sensor in behaving non-human primates, provide a bridge to deeper understanding of the intracellular neural mechanisms of visual information processing.

## Acknowledgments

The authors would like to thank Loren L. Looger for the early provision of AAV-iGluSnFR and the Peking University Laboratory Animal Center for excellent animal care. This work was supported by National Natural Science Foundation of China (grant no. 31730109), National Basic Research Program of China (grant no. 2017YFA0105201), National Natural Science Foundation of China Outstanding Young Researcher Award (grant no. 30525016) and a Project 985 grant of Peking University, Beijing Municipal Commission of Science and Technology (grant no. Z151100000915070), and a U.S. National Science Foundation grant to SLM and SMC (1734887).

## Authors contributions

N.J. and S.T. designed the study, performed experiments and analyzed data. Y.L., F.L. and H.J. performed surgical procedures. N.J., S.T., S.L.M and S.M.C contributed to the interpreted results, and wrote the paper. S.T. supervised the project.

## Methods

### Monkey preparation

All experimental protocols followed the Guide of Institutional Animal Care and Use Committee (IACUC) of Peking University Laboratory Animal Center, and approved by the Peking University Animal Care and Use Committee (LSC-TangSM-5). Rhesus monkeys (*Macaca mulatta*) were purchased from Beijing Prima Biotech, Inc. and housed at Peking University Laboratory Animal Center. Three healthy adult male monkeys (aged 3-4 years and weighing 4-6 kg) were used in this study.

We performed two sequential sterile surgeries for each animal under general anesthesia. In the first surgery, a 20 mm craniotomy was created on the skull over V1 and the dura was then opened. A round cover glass (6 mm in diameter) with a small pore (0.3 mm) was used for targeting the injection pipette and stabilizing cortical surface during injection procedure. The injection pipette was made from a quartz pipette (QF100-70-7.5, Sutter Instrument, USA) and pulled with a 15-20 μm tip using a laser-based pipette puller (P-2000, Sutter Instrument, USA). Through the pipette, 120 to 150 nl of AAV1.hSynap.SF-iGluSnFR.A184S.WPRE.SV40 (HHMI Janelia Research Campus and Addgene, titer 2.7e13 GC/ml) were pressure-injected at a depth of approximately 350 μm. After AAV injections, we inserted a GORE membrane (22 mm in outer diameter) under the dura, sutured the dura, installed back the removed skull bone with titanium pieces and sutured the scalp. The animal was then sent back to its cage for recovery and was administered with Ceftriaxone sodium antibiotic (Youcare Pharmaceutical Group Co. Ltd., China) for one week. We performed a second surgery 45 days later to implant the head posts and imaging window. Three head posts in total were implanted on the animal’s skull, with two of them on the forehead of the skull and one on the back. A T-shaped steel frame was fixed to these head posts for head stabilization during training and experiments. Also in this surgery, the skull was re-opened and subjacent dura was cut off (diameter 16 mm) to exposure the brain cortex. A glass cover-slip (19 mm in diameter and 0.17 mm in thickness) with a titanium ring (12 mm in outer diameter and 10 mm in inner diameter) glued on it was inserted and gently pushed down onto the cortical surface. The titanium ring was glued with dura and skull using dental acrylic to establish an imaging chamber. The whole chamber, reinforced by more thick dental acrylic, was then covered by a steel shell to protect the cover-slip when the animal was returned to the home cage. More relative detailed protocols could be found in a previous study^24^.

### Behavioral task

After ten days recovery from the second surgery, we trained each monkey to sit in a primate chair with its head restrained while performing a fixation task, in which monkeys needed to keep fixation on a small white spot (0.1°) within a window of 2° for over 2.5 seconds to get a juice reward. Eye position was monitored with an infrared eye-tracking system (ISCAN ETL-200, ISCAN Inc. USA) at 120 Hz. Whenever the eye fixated within a scope of 2° of the center white point, a trial started.

### Visual stimuli

Visual stimuli were generated using a ViSaGe system (Cambridge Research Systems) and presented on a 17-inch LCD monitor (Acer V173, 80Hz refresh rate and 1280 × 960-pixel resolution) positioned 45 cm away from the animal’s eyes.

A small drifting grating (full contrast square waves, 4 cycle/degree spatial frequency, 3 cycle/second temporal frequency, 0.2° in diameter) was first used to estimate the RF position of each recorded neuron. For synaptic input tuning measurements (Fig. 1), a set of 81 stimuli were used, which contained 72 drifting square-wave gratings (12 orientations, 2 moving directions at 3 spatial frequencies of 1, 2 and 4 cycle/degree) and 9 color patches (red, orange, yellow, green, cyan, blue, purple, white and black) 1° in size. Direction of motion was not considered in this study and responses across opposing drift directions were averaged. Visual stimuli were displayed in random order. We tested both equiluminant colors (8.2 candela m^-2^) and full saturated colors and found that the results did not vary significantly (Supplementary Fig. 3). Each stimulus was presented for 0.5 s, with an inter-stimulus interval of 0.5 s and repeated 20 times. For receptive field mapping (Fig. 3 and 4), a set of white/black dot stimuli (0.2-0.3° squares in 5 by 5 position array for basal dendritic inputs, and 0.3-0.4° in 7 by 7 position array for apical dendritic inputs) were used. Each condition was repeated 40 times during RF mapping. To precisely measure the time information of synaptic inputs in experiments with high imaging speed, 4-6 preferred stimuli were first chosen for each dendritic branch according to previous functional characterizing experiments. Each stimulus was presented for 100 ms with an inter-trial interval of 100 ms and repeated for approximately 400 times.

### Two-photon imaging

We performed *in vivo* two-photon imaging using a Prairie Ultima IV two-photon microscope (Bruker Nano, Inc., formerly Prairie Technologies) and a Ti: Sapphire laser (Mai Tai eHP, Spectra Physics). A 25× objective lens (1.1 N.A., Olympus) with zoom 4× scanning model was used for high resolution dendritic imaging, which covered a field of view of 136μm × 136μm. Fast resonant scanning mode (512 by 512-pixel resolution, up to 30 frame/second and averaging every 4 frames) was used in all functional characterization experiments. An even faster scanning mode (64 by 64-pixel at 226 frame/second) while zoomed 8× achieved high temporal resolution imaging (without frame averaging, Fig. 4f). To obtain the orientation map, we imaged with a 16× objective lens (0.8-N.A., Nikon) under zoom 1×, covering a field of view of 850μm × 850μm.

### Imaging data analysis

All of the imaging data were analyzed with customized MATLAB codes (Ver. 9.5.0 R2018b, The MathWorks, Natick, MA). To correct the image shifts caused by the relative displacement between the objective and the cortex during imaging process, we first determined a template image by averaging 1000 frames in the middle of each imaging session and then realigned each frame in this session to the template image using a normalized cross-correlation-based translation algorithm^24^. To obtain stimulus-evoked fluorescence changes in the field of view, we subtracted the averaged frame recorded during the stimulus OFF period (4 frames before stimulus onset) from averaged frame from the stimulus ON period (4 frames during stimulus onset) to generate the differential images for each stimulus condition. By using a band-pass Gaussian filter (5 pixels and 50 pixels, 2 orders, respectively), these differential images were spatially filtered and from which, ROIs were identified as individual connected domains consisting of more than 30 pixels with a threshold of 2.5 STD above the differential images. We assumed ROIs as synaptic inputs on neuronal dendrites if they had a 50% intersection with the dendritic domain. To get the morphology of each dendritic branch, we first collected an averaged image throughout each experiment and then filtered this image with a band-pass filter (5 pixels and 50 pixels, 2 orders, respectively). With this processed image, dendritic domain was identified with PS (Adobe Photoshop CC Ver. 19.1.6), and the contour of the dendrites determined using software's magic stick tool (tolerance ranging from 10 to 50 in different conditions to optimize fit). Responses of synaptic inputs were computed as the ratio of fluorescence change (∆*F*/*F*_0_), where ∆*F* = *F* − *F*_0_ and *F*_0_ is the baseline fluorescence intensity during blank screen before stimulus onset in each trial, while *F* is the intensity during stimulus presentation.

### Orientation-selective and color-selective inputs

To collect orientation-selective inputs, the signal-to-noise-ratio (SNR) of single synaptic inputs was defined as the ratio of fluorescence variance between averaged ON and OFF frames of 12 orientations under preferred spatial frequency:

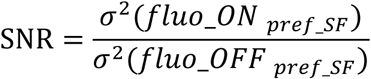

in which *fluo_ON* and *fluo_OFF* are 12 mean fluorescence intensities of target ROI across trials for each orientation condition. Synaptic inputs whose SNR value did not exceed 1 were not taken into the following analysis. Then, we conducted a balanced one-way analysis of variance (ANOVA) on single trial response values of 12 orientations under preferred spatial frequency and set a criteria of p < 0.05 to obtain inputs with significant difference in response strength among 12 orientations. Furthermore, to measure orientation tuning, tuning curves with step size of 15° were fit with circular Gaussian function^28^:

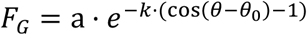

where a is amplitude, *θ*_0_ is the maximum angles and *k* is width parameter (Supplementary Fig. 1). Based on the fitted curves, the orientation selectivity index (OSI)^29,30^ of individual synaptic input was calculated as:

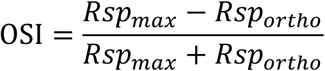

in which *Rsp_max_* is the maximum response strength in optimal orientation and *Rsp_ortho_* corresponded to response value in orientation orthogonal to it (optimal orientation ±90°). Synaptic inputs with OSI exceeding 0.3 were then assigned as orientation-selective inputs. Same SNR index and p-value of ANOVA were used to pick out color-selective inputs.

### Spatial frequency tuning

For each orientation-selective inputs, responses to 3 different spatial frequencies were fit using the Gaussian function (Supplementary Fig. 1). The preferred spatial frequency for each input was the one corresponding to the maximum of fitted tuning curve.

### Orientation map

To demonstrate the spatial organization of orientation preference for cortical areas that we imaged, we first generated raw pseudo-color orientation maps to check data quality manually. Differential images in 12 orientations by averaging 3 spatial frequencies were combined in a raw orientation map, in which color denoted orientation preference and brightness represented relative response intensity. To acquire pinwheel-like orientation maps, the differential images for each orientation were first spatially filtered with a 166-μm (corresponding to 100 pixels) low-pass Gaussian filter. Orientation preference of each pixel on the orientation map was then deduced from the vector sum of its response intensities under 12 orientations^31^:

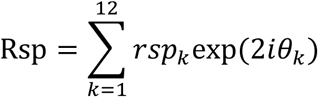

where *θ*_*k*_ were orientations in radians and *rsp*_*k*_ were response intensities. Each single pixel in the orientation map was then assigned to one of 12 bins according to its orientation preference, marked by different colors (Fig. 2h).

### RF map

Individual white (125 candela m^-2^) and black (<0.1 candela m^-2^) squares were presented randomly at one of 25 (5 by 5 matrix) or 49 (7 by 7 matrix) positions covering a total area ranging from 0.8 × 0.8° to 2.6 × 2.6° depending on retinotopic eccentricity of target imaging area. Responses to each single square stimulus were recorded and used to define the ON and OFF subfields of spatial RFs. The 5 × 5 or 7 × 7 response matrix was zoomed in 60 or 40 times respectively and spatially filtered with a Gaussian filter (40 or 26 pixels). Next, a colormap changing from blue to green to red was used to denote ON/OFF signals (Fig. 3 and 4). ON receptive field size was defined as the size of areas in ON RF map with response value higher than a half of maximum response intensity, same for OFF receptive field size. The overall RF size was then defined as the mean value of ON and OFF receptive field size.

### Local clustering

Distance-dependent correlations of individual response properties were calculated along each single dendritic branches. Synaptic inputs on same single dendritic branch were first combined into input pairs (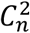 combinations in total), then feature correlation and dendritic distance of each input pair were computed. These input pairs were later allocated into different groups according to their dendritic distance in 2μm steps forming a distance-dependent feature correlation plots (Fig. 2f). We fit these scatter plots with a negative exponential function^12^:

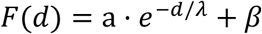

in which a is the maximum correlation value, λ is the spatial length constant and β is the correlation baseline. As a control, synaptic input positions were randomly shuffled and corresponding distance-dependent correlation was calculated for 1000 times. To obtain orientation tuning correlation coefficients, responses to 12 orientations under preferred spatial frequency among orientation-selective inputs were used. For spatial frequency, its tuning was fit as described above and evenly interpolated with 11 data points, with which spatial frequency tuning correlations were computed. Color tuning correlations were calculated using responses to 9 colors among color-selective inputs. Pixel-to-pixel correlations of RF^11^ (5 × 5 or 7 × 7 pixel array according to stimuli set) among orientation-selective inputs were used as a measure of RF similarity.

### Mix of orientation-selective and color-selective inputs

A receiver operating characteristic (ROC) curve^32^ was recruited for analyzing the spatial crossing of orientation-selective and color-selective inputs. Along one specific direction of each dendritic branch, each pixel on dendrite was either located on color-selective ROIs (True positive) or orientation-selective ROIs (False positive), or neither (True negative and False negative). In ROC curves for each single neurons (Supplementary Fig. 3), the horizontal axis represented the ratio of cumulative orientation input area on current point (each point in red line) to overall orientation input area, and the vertical axis marked the corresponding ratio of color inputs. The closer that the ROC curve was to the unity line, the more interlaced the orientation-selective inputs were with the color-selective inputs. Any obvious spatial clustering of orientation-selective or color-selective inputs on dendrites would lead to a deviation of ROC curve away from the unity line, causing the area under curve (AUC) deviating from the value of 0.5.

### High-resolution temporal tuning

Several preferred stimuli for each dendritic branch were used in experiments with high imaging speed up to 226 Hz. For each synaptic input, mean fluorescence intensity through 10 frames prior to stimulus onset across trials was assigned as fluorescence baseline. To get an averaged time curve of overall inputs on one dendritic branch, temporal tunings of each synaptic input under its optimal stimulus with maximum response intensity were weighted summed by each ROI’s area size. Time curves of apical and basal dendrites were then normalized according to the peak response value and then fit with a fourth-order Fourier function respectively (Fig. 4f). The time point corresponding to the half-height of each fitted time curve was collected for comparison of temporal information.

### Statistics

Sampling size of neuron and synaptic input number are similar to other relative studies in this field, and exact sample sizes were described in the text. The degree of correlation was calculated as Pearson’s linear correlation coefficient in this study unless otherwise specified. We used non-parametric Kruskal-Wallis test to compare response intensities of overall synaptic inputs on different neurons with a following Multiple Comparison test (Fig. 1). ROC analysis was conducted to illustrate the spatial interlacing relationship of orientation-selective and color-selective inputs. The linear dependence between two distinct variables was characterized by P-value or R-square of linear regression analysis (Fig. 2–4 and Supplementary Fig. 4-7). The experimenter was blind to each neuron’s location when measuring orientation column structure. To compare functional properties of synaptic inputs on apical and basal dendrites of same single neuron (Fig. 4), we employed the Paired Sample T-test analysis. No estimates of statistical power were executed before experiments.

### Code and data availability

Data and custom code are available upon reasonable request.

