## Supplementary material for "Spatiotemporal functional organization of excitatory synaptic inputs onto macaque V1 neurons": Supplemantal Information

### Spplementary information

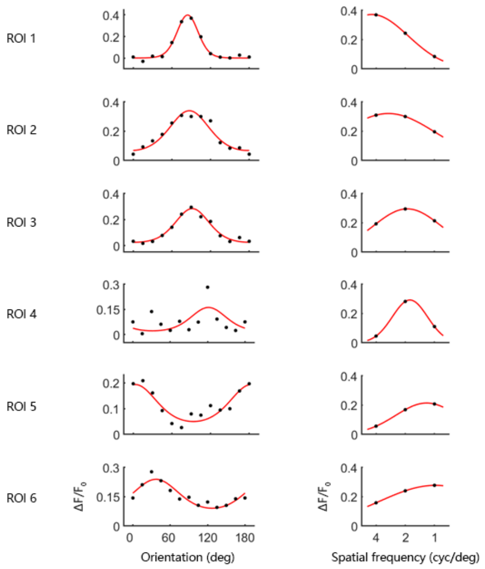


**Supplementary Fig. 1 Examples of orientation and SF tuning of dendritic inputs.** 6 ROIs from the first neuron in **Fig. 2**. Left column, responses to different orientations under optimal SF. Black dots, raw data points; red curves, circular Gaussian curve fits. Right, responses to 3 spatial frequencies under optimal orientation. Black dots, raw data points; red curves, circular Gaussian fits.

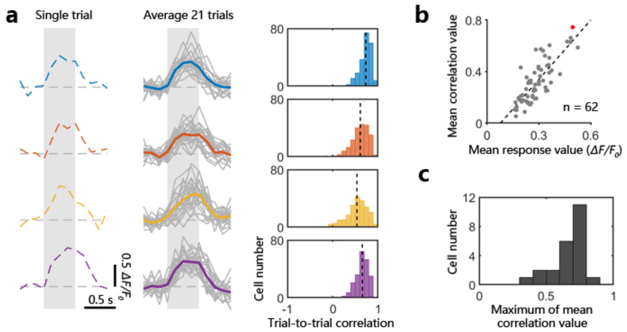


**Supplementary Fig. 2 Single-trial repeatability.** **a**, Single-trial fluorescence transients from the corresponding ROIs in **Fig. 1a** under each ROI's preferred stimuli. Left, single trials; middle, average (colored) of 21 individual trials (gray); right, distribution of trial-to-trial correlations of time courses (blue, $0.73 \pm 0.12$; orange, $0.61 \pm 0.18$; yellow, $0.54 \pm 0.22$; purple, $0.65 \pm 0.13$; mean$\pm$SD). **b**, Mean trial-to-trial correlation value versus mean response value for all 62 ROIs on the target neuron dendrites. Red dot, the ROI with the highest mean correlation value; black dash line, best-fit linear regression line ($p = 6 \times 10^{-15}$). **c**, Statistics of the highest mean correlation value for each neuron ($n = 23$).

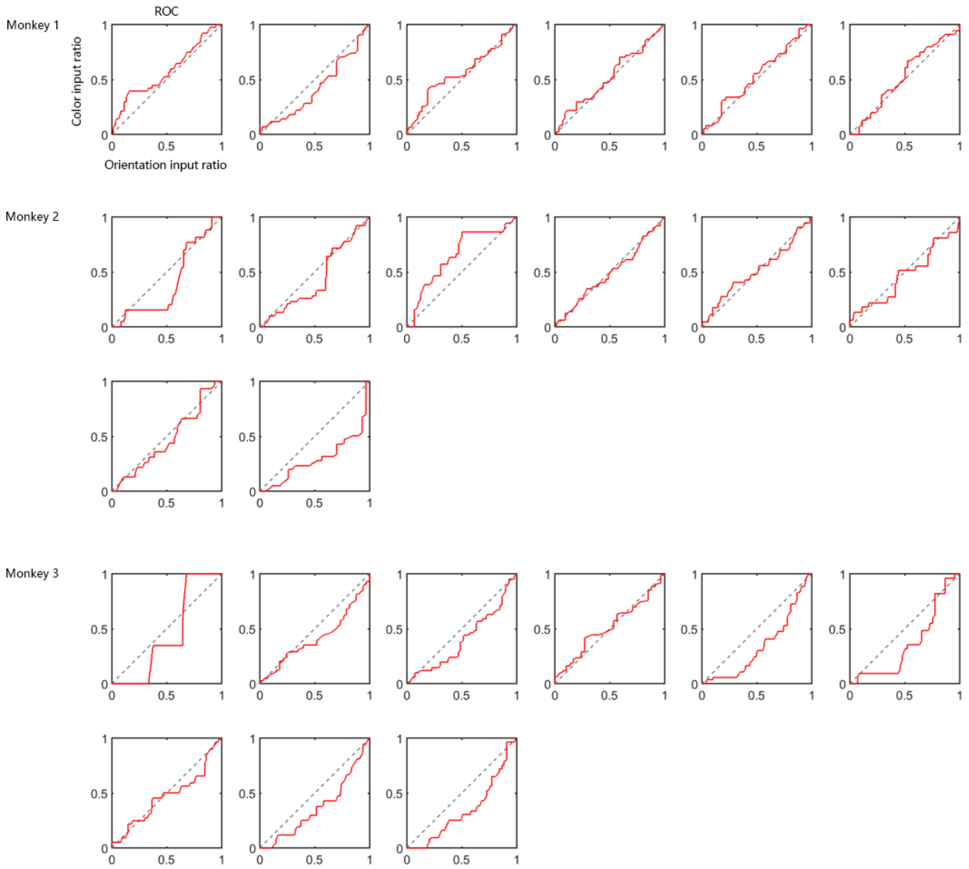


**Supplementary Fig. 3 Highly hybrid distribution of orientation-selective and color-selective inputs on neuronal dendrites.** ROC analysis of synaptic input type distribution along dendrites in each neuron ($\mathrm{AUC} = 0.46 \pm 0.08$, area under curve, $\mathrm{mean} \pm \mathrm{STD}$ among 23 neurons from 3 monkeys in total). More details in Methods.

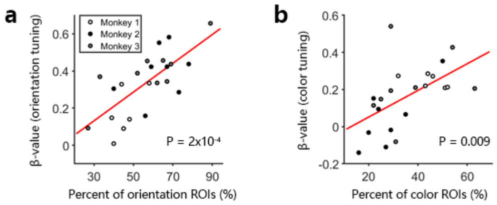


**Supplementary Fig. 4 Correlation between functional properties and composition of synaptic inputs. a**, Relationship between the baseline orientation correlation between ROIs (β-value) versus the ratio of orientation-selective ROIs for each neuron ($N = 23$). Red line, linear fit ($y = kx + b$, $k = 0.0078 \pm 0.0035$, $b = -0.1014 \pm 0.2054$; 95% condifence intervals; $p = 2 \times 10^{-4}$); white dots, neurons from Monkey 1; black, Monkey 2; grey, Monkey 3. **b**, Same as panel **a**, for color tuning correlation as a function of color-selective ROIs percentage ($N = 22$; $k = 0.0072 \pm 0.0051$, $b = -0.0924 \pm 0.1947$; $p = 0.009$). The high correlations suggest that the proportion of inputs of a given type specifies the baseline function of the neuron.

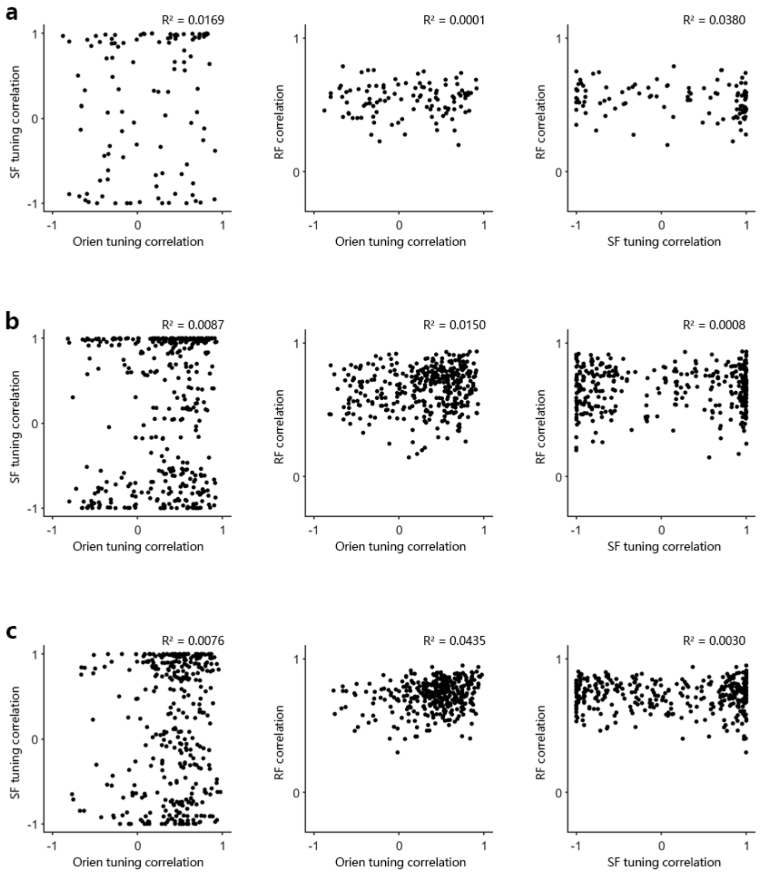


**Supplementary Fig. 5 Inter-input property relationship among various feature dimensions.** Same layout as Fig. 3 c-e, inter-input properties of neurons in Fig. 2i were presented here as **a-c**.

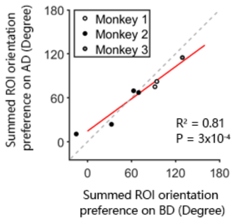


**Supplementary Fig. 6** Orientation preference of summed orientation-selective ROIs on AD versus that on BD for single neurons ($n = 7$, neurons with less than 10 orientation-selective ROIs on AD were discarded). Red line, regression line ($p = 3 \times 10^{-4}$); grey dashed line, unity line. $R^2 = 0.81$ when fitted to the unity line.

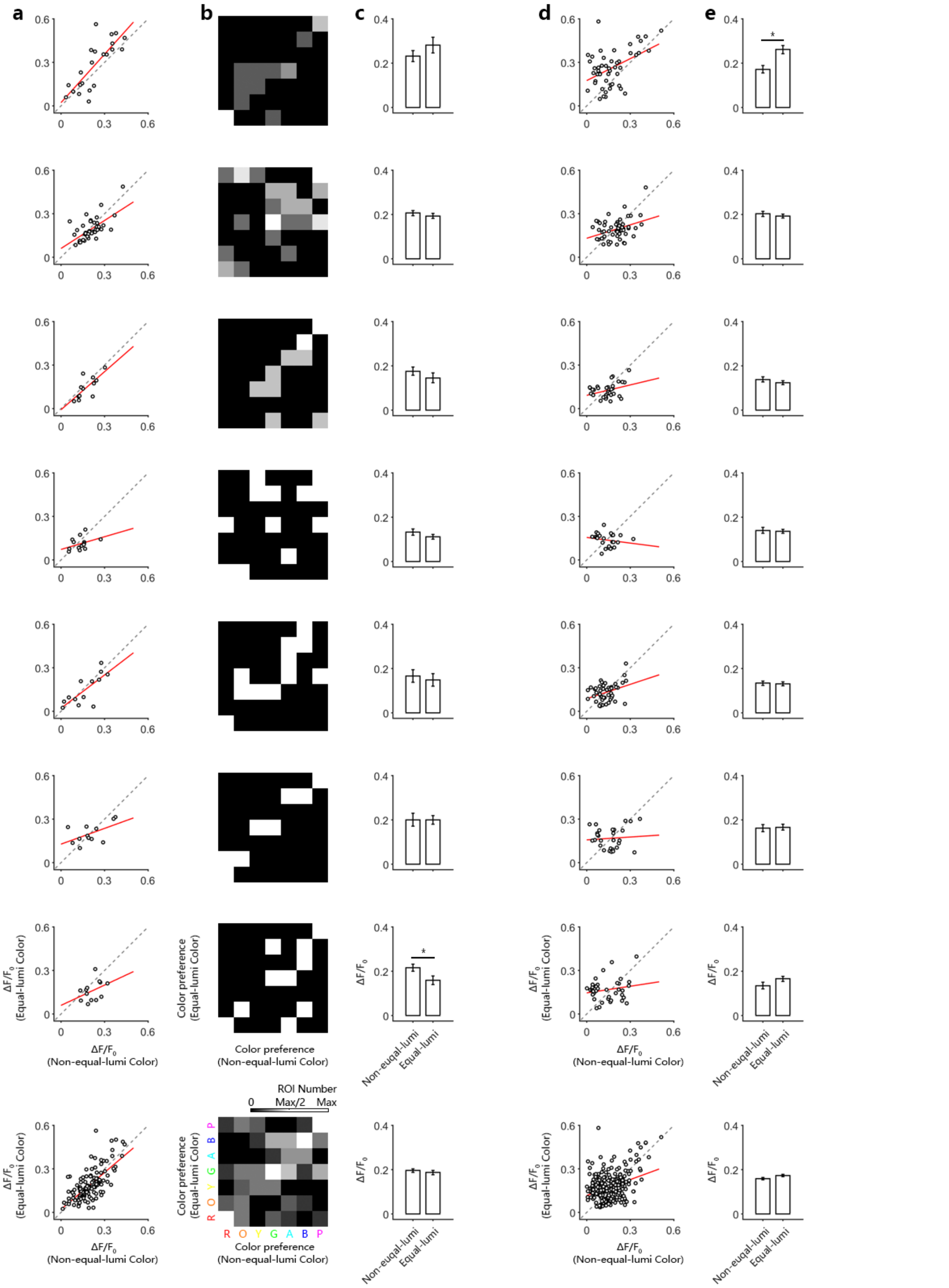


**Supplementary Fig. 7 Comparison between non-equiluminant and equiluminant chromatic stimulus evoked responses. a**, Peak response intensities of ROIs to equiluminant color stimuli versus non-equiluminant (but colored) stimuli. Red line, linear regression. 7 neurons in total from Monkey 3 were recorded. Bottom, statistics for all ROIs from all 7 neurons. **b**, Color preference of each ROI among equal-luminance colors versus non-equal-luminance colors. Brightness denotes ROI proportion, referring to colorbar above the bottom figure. **c**, Averaged response intensity of all ROIs under non-equiluminant versus equiluminant chromatic stimuli. ROIs in **a**-**c** were collected only according to non-equiluminant color stimuli. Asterisk, significant difference, by Wilcoxon rank sum test; $p = 0.11$ when combining all ROIs. **d**, Non-equiluminant and equiluminant chromatic stimuli response strengths within ROIs that were tested with both. **e**, Same as **c**, statistics of averaged response intensities; $p = 0.07$ when combining all ROIs.

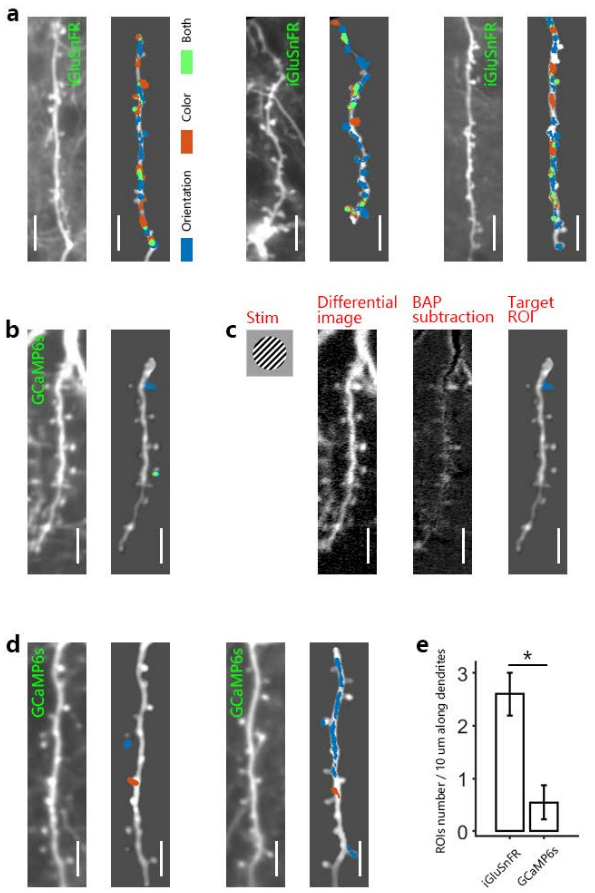


**Supplementary Fig. 8 Comparison between iGluSnFR and GCaMP6s recorded dendritic activities in macaques. a**, Dendritic shafts from distinct neurons expressing SF-iGluSnFR.A184S in superficial layer of V1 and visual stimuli evoked ROIs on them. Scale bar, 10 µm. **b**, Dendritic shaft of one neuron expressing GCaMP6s and corresponding ROIs. **c**, A subtraction procedure to minimize contamination from back-propagation action potentials (BAPs). Leftmost, one visual stimulus; middle left, the differential image of responses under this visual stimulus; middle right, the response image after subtracting BAPs; rightmost, the target ROI selected. **d**, Another two samples of dendritic shafts from neurons expressing GCaMP6s and corresponding ROIs. **e**, Density comparison of recorded ROIs on iGluSnFR and GCaMP6s expressing neural dendrites.
